# Rational Discovery of Dual-Action Multi-Target Kinase Inhibitor for Precision Anti-Cancer Therapy Using Structural Systems Pharmacology

**DOI:** 10.1101/465054

**Authors:** Hansaim Lim, Di He, Yue Qiu, Patrycja Krawczuk, Xiaoru Sun, Lei Xie

## Abstract

Although remarkable progresses have been made in the cancer treatment, existing anti-cancer drugs are associated with increasing risk of heart failure, variable drug response, and acquired drug resistance. To address these challenges, for the first time, we develop a novel genome-scale multi-target screening platform 3D-REMAP that integrates data from structural genomics and chemical genomics as well as synthesize methods from structural bioinformatics, biophysics, and machine learning. 3D-REMAP enables us to discover marked drugs for dual-action agents that can both reduce the risk of heart failure and present anti-cancer activity. 3D-REMAP predicts that levosimendan, a drug for heart failure, inhibits serine/threonine-protein kinase RIOK1 and other kinases. Subsequent experiments confirm this prediction, and suggest that levosimendan is active against multiple cancers, notably lymphoma, through the direct inhibition of RIOK1 and RNA processing pathway. We further develop machine learning models to identify cancer cell-lines and patients that may respond to levosimendan. Our findings suggest that levosimendan can be a promising novel lead compound for the development of safe and effective multi-targeted cancer therapy, and demonstrate the potential of genome-wide multi-target screening in designing polypharmacology and drug repurposing for precision medicine.

**Author Summary:** Multi-target drug design (a.k.a targeted polypharmacology) has emerged as a new strategy for discovering novel therapeutics that can enhance therapeutic efficacy and overcome drug resistance in tackling multi-genic diseases such as cancer. However, it is extremely challenging for conventional computational tools that are either receptor-based or ligand-based to screen compounds for selectively targeting multiple receptors. Existing multi-target drug design mainly focuses on compound screening against receptors within the same gene family but not across different gene families. Here, we develop a new computational tool 3D-REMAP that enables us to identify chemical-protein interactions across fold space on a genome scale. The genome-scale chemical-protein interaction network allows us to discover dual-action drugs that can bind to two types of targets simultaneously, one for mitigating side effect and another for enhancing the therapeutic effect. Using 3D-REMAP, we predict and subsequently experiments validate that levosimendan, a drug for heart failure, is active against multiple cancers, notably, lymphoma. This study demonstrates the potential of genome-wide multi-target screening in designing polypharmacology and drug repurposing for precision medicine.

## Introduction

In spite of the tremendous success of kinase-targeting drug and immunotherapy in the treatment of cancers, the development of anti-cancer targeted therapeutics faces three challenges: serious side effects, especially, cardiotoxicity, variable drug responses, and acquired drug resistance. For example, tyrosine kinase-targeting drugs are shown to be associated with a higher risk of onset heart failure in adult cancer patients[1]. Recently, fatal heart failures have been reported in patients treated with immune checkpoint inhibitors[2].

The clinical responses of immune checkpoint inhibitors strongly depend on the interplay of microenvironment and global immunity[3]. As a result, only a small portion of cancer patients respond to immunotherapy. Similarly, the efficacy of kinase-targeting cancer therapeutics depends on the catalytic activity of the targeted kinase[4]. Existing kinase drugs mainly involve targeting a kinase that is directly activated by mutation. Due to the high heterogeneity of cancers, many patients may not harbor the activated mutation of targeted kinases. Thus, it is still urgently needed to target other cancer mechanisms as well as identify patients associated with such aberrations for effective and precision anti-cancer therapy.

In addition to primary resistance to immunotherapy and kinase-targeted drugs, adaptive and acquired resistance to anti-cancer therapies inevitably emerges[5, 6], even in cancer patients who initially respond to the therapeutics. Acquired resistance is mediated by multiple mechanisms such as modification of targeted kinases, activation of bypass signaling pathways, or histological transformation. Drug combinations that can target multiple cancer pathways has been actively pursued to combat the drug resistance in the treatment of cancer[7].

Put together, multi-targeted therapy through either drug combination or a single polypharmacological agent that targets multiple disease-associated mechanisms could be an effective and efficient strategy to mitigate side effect, enhance therapeutic efficacy, and check drug resistance in the cancer treatment. Although the multi-targeted therapy has been discovered to either mitigate side effect[8] or enhance therapeutic efficacy alone[9], few studies have been reported to design molecules as a dual-action agent that can achieve both of objectives at the same time. In this paper, we for the first time develop a novel genome-wide multi-target screening platform and apply it to discover marked drugs that could be the dual-action agent. Our method is based on the premise that a drug often interacts with unintended targets, i.e., off-targets. These off-targets may cause unwanted side effects (i.e., anti-target) or function as therapeutic targets for another disease. The objective of our method is to identify such targets and associated marked drugs that can selectively interact with multiple targets. These off-targets could collectively reduce side effect and enhance therapeutic effect.

Selective promiscuity has been actively pursued in kinase inhibitor design[10]. However, current efforts in multi-target screening only consider the binding promiscuity within the kinase family. The dual-action drug design strategy needs to extend the concept of selective promiscuity across gene families. It is challenging to identify the genome-wide drug-target interactions[11]. In this paper, we develop a novel platform 3D-REMAP to screening off-targets of marked drugs on a genome scale. The 3D-REMAP integrates diverse chemical genomics, structural genomics, and functional genomics data as well as combines various computational tools from bioinformatics, chemoinformatics, biophysics, and machine learning. As a result, the 3D-REMAP partially overcomes the limitations of each individual data set and method. To our knowledge, it is the first time that the multi-target screening has been applied to the precision drug repurposing.

Using the 3D-REMAP, we have made several innovative discoveries in this paper. First, we discover that levosimendan, a drug for heart failure, is a novel inhibitor of serine/threonine-protein kinase RIOK1 and a number of other kinases. Second, we uncover that RIOK1 and its associated RNA processing pathway is an effective novel target for multiple types of cancers, especially, lymphoma. Different from existing targeted kinases that harbor the activated mutation, the aberration of RIOK1 is mainly associated with its overexpression. Thus targeting RIOK1 may provide new opportunities in the cancer treatment. Third, we suggest that levosimendan can be a novel lead compound for developing a more safe and effective anti-cancer therapy to overcome the cardiotoxicity and the drug resistance of existing kinase-targeting drugs, either as a single polypharmacological agent or a component in a drug combination. Our findings may have significant implications in anti-cancer drug discovery, and demonstrate the potential of the genome-wide multi-target screening in precision drug discovery.

## Results

### 1. Rational polypharmacology strategy for discovering dual-action multi-target agents

The rationale of our multi-target drug screening strategy is that the serious side effect caused by therapeutic target(s) (i.e. on-target effect) or anti-target(s) (i.e. off-target effect) of a drug can be mitigated by its or another drug’s interaction with an off-target, which is against the side effect, as shown in Figure 1A. The drug-off-target interaction can come from a single polypharmacological agent or multiple components in a drug combination. In this study, we focus on searching for marked drugs that may act as a dual-action agent that can mitigate the cardiotoxicity of anti-cancer therapy, at the same time, present anti-cancer potency. Contemporary anti-cancer therapeutics, such as tyrosine kinase inhibitors, anthracycline chemotherapy, and immunotherapy, are all associated with the cardiotoxicity[12].

**Figure 1.**
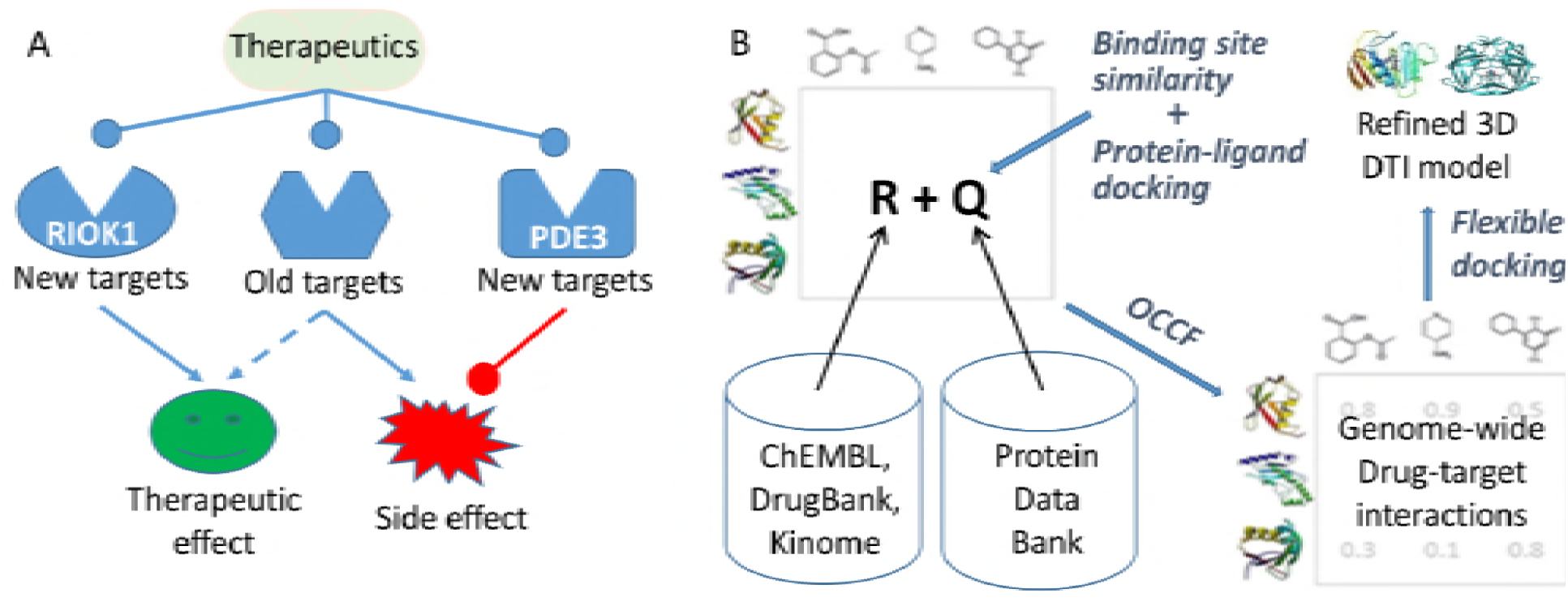
(A) A rational polypharmacology strategy to screen drugs that can both enhance therapeutic effect and mitigate side effect. (B) Schema of 3D-REMAP, a multi-target screening platform that integrates structural genomics and chemical genomics data and combines tools from bioinformatics, chemoinformatics, biophysics, and machine learning. *R* and *Q* denote observed protein-chemical interactions in chemical genomics databases, and predicted protein-chemical interactions from ligand binding site similarity coupled with protein-ligand docking, respectively. These two matrices are the input for the machine learning algorithm One-Class Collaborative Filtering (OCCF) to predict genome-wide drug-target interactions. See method section for details. DTI: drug-target interaction.

### 2. Hypothesis generation using 3D-REMAP, a genome-scale multi-target screening platform

The rational discovery of the dual-action therapeutic agent was achieved using a novel genome-scale multi-target screening platform 3D-REMAP, as illustrated in Figure 1B. The 3D-REMAP applied here integrated diverse data from structural genomics and chemical genomics, as well as synthesized tools from bioinformatics, chemoinformatics, biophysics, and machine learning. Since each data set alone is biased, incomplete, and noisy, and a single computational or experimental tool has its inherited advantages and limitations, the integrative analysis may provide more comprehensive and reliable results. The 3D-REMAP takes four networks as input: a protein-chemical interaction (PCI) network, which is represented by matrix *R*. Since observed PCI is highly sparse, one of unique features of 3D-REMAP is to apply ligand binding site similarity search and protein-ligand docking to screen off-targets of a given drug across human structural proteome[13-19]. The predicted off-target network (represented by matrix *Q*) fills in the part of unobserved entries in the PCI network. In addition to *R* and *Q*, two other input networks are a chemical-chemical similarity network and a protein-protein similarity network (not shown in Figure 1B). In this work, the observed PCI network is derived from ChEMBL[20] (v23.1), DrugBank[21] (v5.5.10), 2), and four kinome assay data sets[22-25]. The final network includs 1,656,274 unique chemicals, 9,685 unique target proteins, and 650,581 observed interactions. Then a weighted imputed neighborhood regularized One-Class Collaborative Filtering (OCCF) algorithm developed by us[26], which can handle uncertainty and missing values in the observed and putative interactions, is applied to predict the binding profile of the given drug againt the 9,685 targets. Finally, the atomic details of the prioritized drug-target interactions are analyzed for lead optimization using flexible protein-ligand docking.

### 3. Protein kinases are predicted to be the off-target of PDE3 inhibitors

In this study, five drugs (milrinone, anagrelide, amrinone, enoximone, and levosimendan) that were used for the treatment of heart failure were selected as our starting point. The reported primary targets of these drugs include PDE3[27], potassium channel[28], and troponin C[29]. Using PDE3B structure as a template, the potential off-targets of PDE3B were predicted using SMAP software[30-32] and protein-ligand docking[33]. To increase the coverage of proteome, the relatively high confident predicted drug-target interactions from SMAP and protein-ligand docking (SMAP p-value<0.005, and protein-ligand docking score < -7.5) were incorporated into the genome-scale PCI network, and 3D-REMAP was applied to predict drug binding profiles. 3D-REMAP not only is scalable to tens of thousands of protein targets on a genome scale but also takes the uncertainty of observed or inferred drug-target interactions into consideration, and consolidate their consistency or conflict relations in a global context. Thus, it may predict off-targets that are missed by the protein structure-based approach and other chemoinformatics methods[26, 34].

Among those off-targets predicted by 3D-REMAP, we focused on protein kinases, since a significant number of them are known to be associated with cancers. In addition, it is relatively easy to experimentally validate the predicted off-targets across human kinome. Levosimendan, a marketed drug in Europe and Asia for the heart failure, was selected for the initial experimental validations. The top 3 predicted kinase off-targets of levosimendan are CAMK2, RIOK1, and FLT3. Additional top-ranked predictions include RIOK3, MYLK4, LTK, CDK7, CDK8, DYR1B, GSK3A, GSK3B, and MAP3K5. The expectation values of all of these predictions are less than 1.0e-11.

### 4. RIOK1 and several other kinases are the off-targets of levosimendan

KinomeScan^™^ assay across 452 human kinases verified our computational prediction (Supplemental Data S1). RIOK1 is one of the strongest inhibited kinases by levosimendan. The percentage control of RIOK1 is 15.0 and 0.0 under the treatment of 10μM and 100 μM of levosimendan, respectively. RIOK3, the closest homolog of RIOK1, showed the same binding strength as RIOK1. In addition, levosimendan also inhibits a number of other kinases besides RIOK1 and RIOK3. The percentage controls of five kinases: FLT3, MAP2K5, PIP5K1A, GAK, and KIT are less than 30.0 under the treatment of both 10 μM and 100 μM of levosimendan. Their distributions in the kinome tree is shown in Figure 2. Both FLT3 and KIT are tyrosine protein kinases. Their inhibitors have recently been proved for the treatment of Acute Myeloid Leukemia and other types of cancers, but a combination therapy is needed to overcome rapidly emerged drug resistance, thus improve patients’ drug responses[35, 36]. Other kinases inhibited by levosimendan belong to serine/threonine protein kinase family (RIOK1, RIOK3, GAK, and MAP2K5) and lipid kinase family (PIP5K1A). As a result, levosimendan may modulate different cancer pathways from those by the tyrosine protein kinase inhibitors, and could be a novel lead compound for a new anti-cancer therapy to overcome drug resistance of existing drugs or to target different types of cancers through polypharmacology or drug combination.

**Figure 2.**
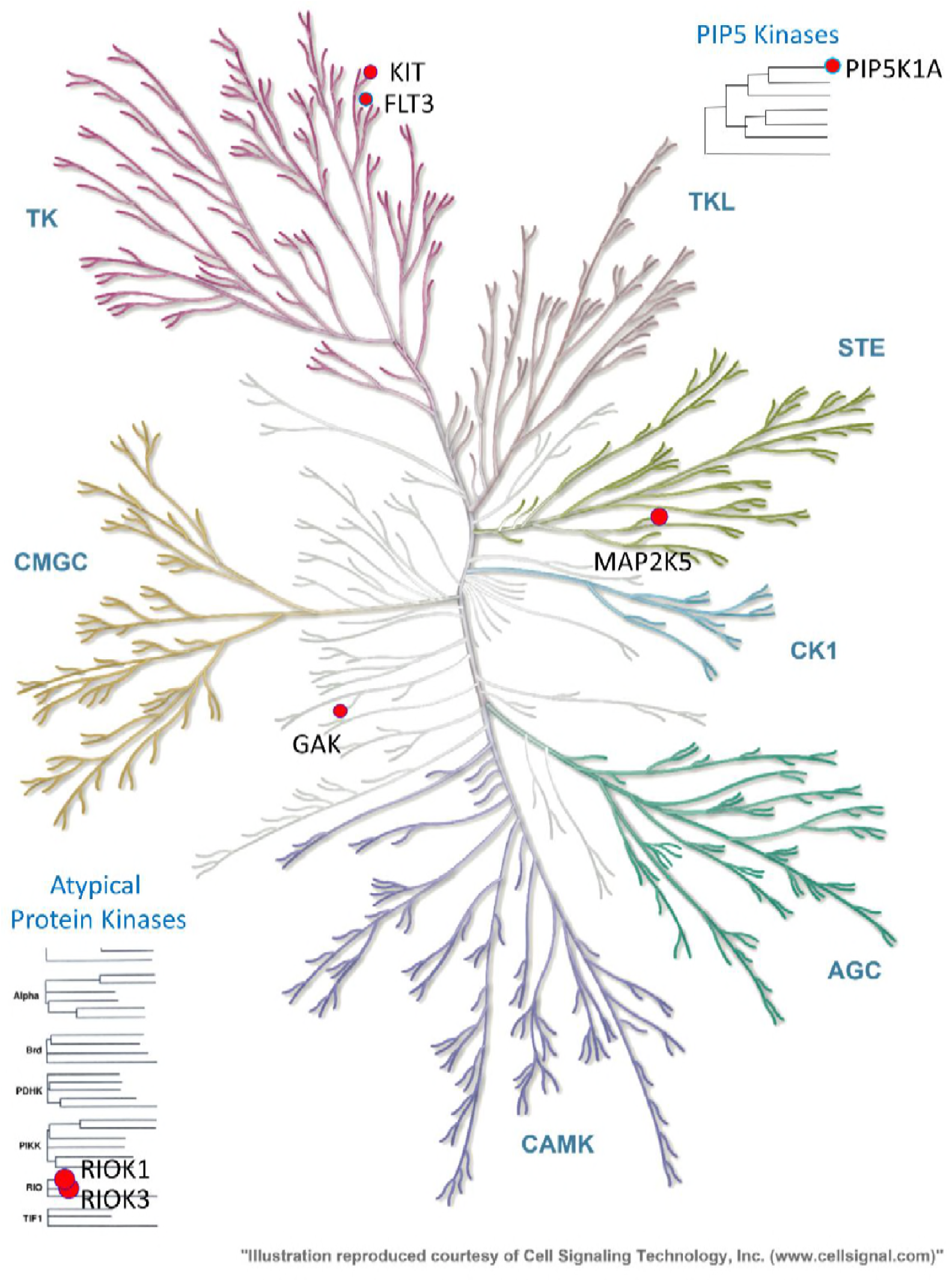
The distribution of kinase off-targets of levosimendan in the human kinome. The off-targets are marked by red circles. The diameter of the circle approximately corresponds to the binding strength.

To understand the molecular details in how levosimendan interacts with RIOK1, we docked levosimendan into the ATP binding site of RIOK1 using AutodockFR, a flexible receptor protein-ligand docking program[37]. The binding pose and interaction pattern between levosimendan and RIOK1 is shown in Figure 3A and 3B, respectively. The orientation of levosimendan is close to that of ADP. Several key interactions between ADP and RIOK1 observed in the crystallized complex structure (PDB id: 4OTP) are conserved in the levosimendan-RIOK1 complex. They include the hydrogen bonds formed by ILE280, SER187, and water molecules as well as Pi-Alkyl interaction formed by VAL194. Such information may facilitate the medicinal chemistry efforts in optimizing levosimendan to be a more potent selective RIOK1 inhibitor. For example, a functional group of hydrogen bond acceptor may be added to the benzene ring to form hydrogen bond interaction with ASP341. The amino acid mutations in the binding site may impact the ligand binding. No amino acid mutations in TCGA[38] and COSMIC[39] are observed in the key residues involved in the binding of levosimendan. Thus, levosimendan may target the aberration of RIOK1 that is associated with overexpression or post-translational modification of protein rather than amino acid mutations, and distinguish itself from existing kinase inhibitors. It is noted that the binding site residues of RIOK1 and RIOK3 are highly conserved. It could be difficult to design ligands that selectively target RIOK1 but not RIOK3.

**Figure 3.**
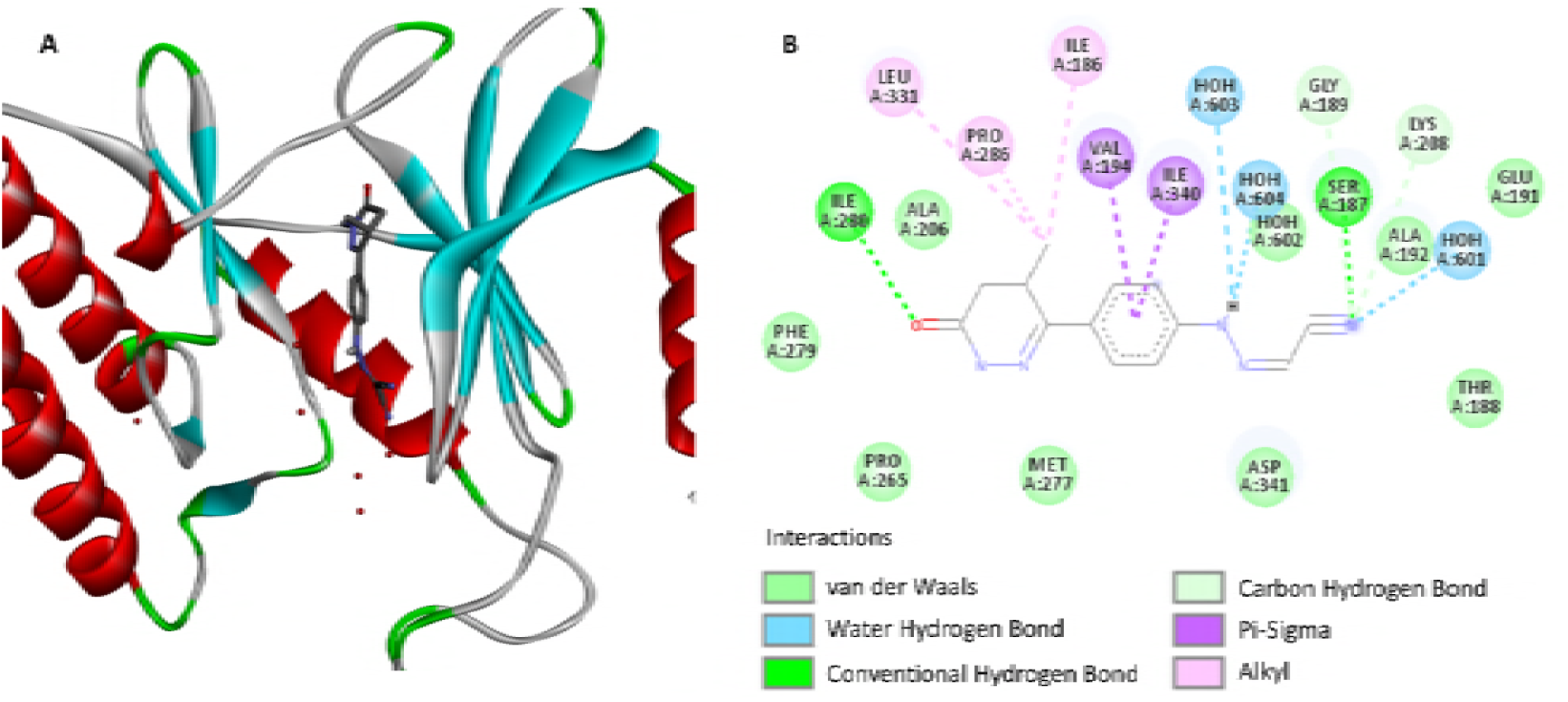
Predicted binding pose (A) and interaction pattern (B) of levosimendan with RIOK1. RIOK1 and levosimendan is represented by ribbon and stick model, respectively.

Comparing the computationally predicted kinase off-targets of levosimendan with the experimentally determined ones across human kinome, we correctly predicted 2 off-targets out of top 3 ranked predictions (RIOK1 and FLT3). Among the top 12 ranked predictions, three predicted off-targets have strong bindings to levosimendan (RIOK1, RIOK3, and FLT3), two have intermediate binding strengths (MYLK4 and CAMK2), and others are false positives (LTK, CDK7, CDK8, DYR1B, GSK3A, GSK3B, and MAP3K5). There are four false negatives (MAP2K5, PIP5K1A, GAK, and KIT). There is no doubt that the performance of 3D-REMAP needs to be further improved. Nevertheless, the successfully predicted off-targets of levosimendan from 3D-REMAP cannot be achieved by conventional protein-ligand docking or ligand-based virtual screening alone (Supplemental Data S2 and S3).

### 5. Levosimendan can inhibit the proliferation of multiple types of cancer cells

The putative anti-cancer activity of levosimendan was tested over 200 cancer cell lines across 19 cancer types (sites of primary tumor), which include bladder, breast, central nervous system, colon, endocrine, eye, female genitourinary system, head and neck, hematopoietic, kidney, liver, lung, pancreas, placenta, prostate, skin, soft tissue, stomach, and testis (Supplemental Data S4). Among them, the EC_50_, IC_50_, and GI_50_ of 17 cell lines were all less than 10.0 μM. Hematopoietic Lymphoma was the most sensitive to levosimendan. 4 out of 17 the most sensitive cell lines belong to the lymphoma cancer type. Notably, the EC_50_, IC_50_, and GI_50_ for SU-DHL-8 cell line are 0.604 μM, 0.604 μM, and 0.512 μM, respectively. Figure 4 shows the drug response curve of the SU-DHL-8 cell line under the levosimendan treatment. The cell count activity area is 5.0, significantly higher than those of other tested cell lines. Other cancer types sensitive to levosimendan include stomach carcinoma, endocrine carcinoma, kidney tumor, colorectal carcinoma, bladder carcinoma, osteosarcoma, melanoma, prostate hyperplasia, and sarcoma. Thus, levosimendan is a promising lead compound for designing polypharmacological agent or drug combination to treat multiple types of cancers, notably, lymphoma.

**Figure 4.**
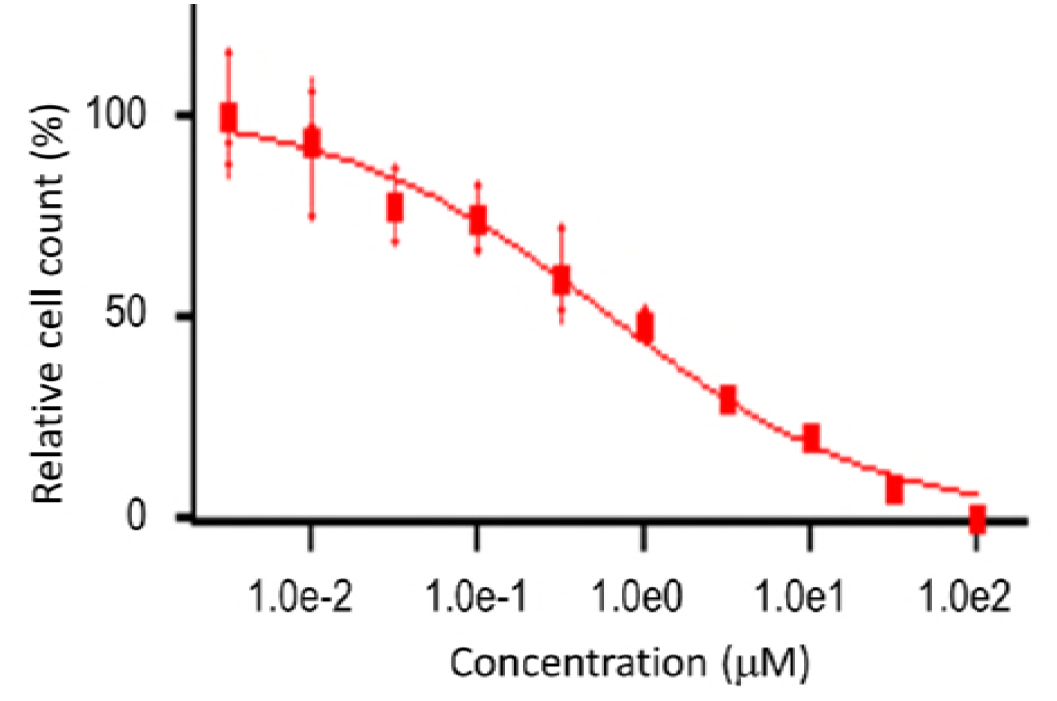
The drug response curve of lymphoma SU-DHL-8 cell line under the treatment of levosimendan.

### 6. Anti-cancer activity of levosimendan may come from its modulation of RNA processing pathway through the inhibition of RIOK1

Differential gene expression profile analysis across 200 cancer cells shows 475 genes may contribute to the cell line sensitivity to levosimendan with *q*-value less than 1.0e-3 (Supplemental Data S5). Gene Set Overrepresentation Analysis suggests that the drug sensitivity genes are significantly enriched with the biological processes of Gene Ontology[40]: rRNA processing, translation, and ribosome biogenesis (FDR<1.0e-3), as shown in Table 1. The only overrepresented KEGG pathway is “ribosome” (FDR=6.62e-40).

**Table 1.** Overrepresented GO biological process terms responsible for the anti-cancer sensitivity of levosimendan.

| **Term** | ***p*-value** | **Fold Enrichment** | **Bonferroni** | **Benjamini** | **FDR** |
| --- | --- | --- | --- | --- | --- |
| GO:0006364<br>rRNA processing | 6.35E-50 | 16.74795907 | 5.68E-47 | 5.68E-47 | 9.89E-47 |
| GO:0006413<br>translational initiation | 1.49E-48 | 22.2853351 | 1.33E-45 | 6.65E-46 | 2.31E-45 |
| GO:0006614<br>SRP-dependent co-translational protein targeting to membrane | 7.32E-47 | 28.24320915 | 6.55E-44 | 1.64E-44 | 1.14E-43 |
| GO:0019083<br>viral transcription | 1.60E-46 | 24.88932806 | 1.43E-43 | 2.86E-44 | 2.49E-43 |
| GO:0006412<br>translation | 3.16E-39 | 12.85456733 | 2.83E-36 | 4.72E-37 | 4.92E-36 |
| GO:0002181<br>cytoplasmic translation | 6.02E-08 | 21.23889328 | 5.39E-05 | 7.70E-06 | 9.38E-05 |
| GO:0042274<br>ribosomal small subunit biogenesis | 7.60E-08 | 29.03754941 | 6.80E-05 | 8.50E-06 | 1.18E-04 |

RIOK1 is overexpressed in the drug sensitive cell lines, and one of the 475 genes. It is established that the primary function of RIOK1 and RIOK3 are involved in the rRNA processing of ribosome[41, 42]. Thus, differential gene expression and gene set overrepresentation analyses are consistent with our computational prediction and kinase binding assay. It is reasonable to hypothesize that the inhibition of RIOK1 by levosimendan is directly responsible for its anti-cancer activity.

Differential expression analysis also unveiled genes that were significantly associated with resistance to levosimendan. The top genes in this set are involved in a wide variety of cellular functions, including signal transduction, mitosis, cytoskeletal regulation, ion transport, and drug metabolism, but no biological processes or pathways are overrepresented.

Different from gene expression profiles, amino acid mutations and copy number variations associated with the sensitivity and resistance of levosimendan were statistically insignificant. This was consistent with the predicted binding pose of levosimendan in RIOK1. Thus, levosimendan may represent a new class of kinase inhibitors that do not depend on targets activated by mutations.

### 7. Pharmacogenomics modeling of anti-cancer activity of Levosimedan

Using the expression values of genes that are responsible for the sensitivity and resistance of levosimendan as features, we applied a novel feature selection method Kernel Conditional Covariance Mininization[43] to develop machine learning models for the prediction of other cancer cell lines or patients that may respond to levosimendan. We developed two models using CCLE[44] and GDSC[45] data separately. The Spearman’s correlation for these two models was 0.7547 and 0.6727, respectively. In the leave-one-out cross-validation, the predicted activity area for the most active SU-DHL-8 cell line was around 2.0. Thus the predicted activity area of 2.0 was used as a threshold for the prediction. Supplemental Data S6 listed top 50 ranked predictions.

Top 30 ranked cell lines that were predicted with consensus by both CCLE and GDSC models included JM1 (B cell lymphoma), NU-DUL-1 (B cell lymphoma), SU-DHL-5 (B cell lymphoma), and ALL-SIL (T-Cell Leukemia). Their predicted active areas were larger than 3.8 and 2.0, respectively. It is clear that B-cell lymphoma dominates the sensitive cell lines to levosimendan.

We further applied the trained CCLE model to predict the cases in TCGA[38] that could respond to levosimendan (Supplemental Data S7). Consistent with the cell line assays and predictive models, the top 100 ranked cases are overrepresented by B-cell lymphoma (TCGA-DLBC) with the *p*-value of 1.05e-2, as shown in Figure 5. The predicted activity areas for the cases of B-cell lymphoma span a broad range from less than 0.2 to larger than 2.0, suggesting that only a portion of B-cell lymphoma patients may respond to the treatment of levosimendan. Except B-cell lymphoma, other TCGA projects are not significantly overrepresented in the top 100 ranked predictions. However, a number of cases in multiple cancer types have the predicted active area larger than 2.0, and fall into the cancer types that respond to levosimendan in the cell line assay. Notably, they include stomach carcinoma (TCGA-STAD), kidney renal clear cell carcinoma (TCGA-KIRC), prostate adenocarcinoma (TCGA-PRAD), colon adenocarcinoma (TCGA-COAD), bladder urothelial carcinoma (TCGA-BLCA), and skin cutaneous melanoma (TCGA-SKCM). Thus, some patients diagnosed with these cancers may also benefit from the treatment of levosimendan. Due to the heterogeneity of cancers, it is necessary to develop an accurate pharmacogenomics model for the development of levosimendan as a precision anti-cancer therapy.

**Figure 5.**
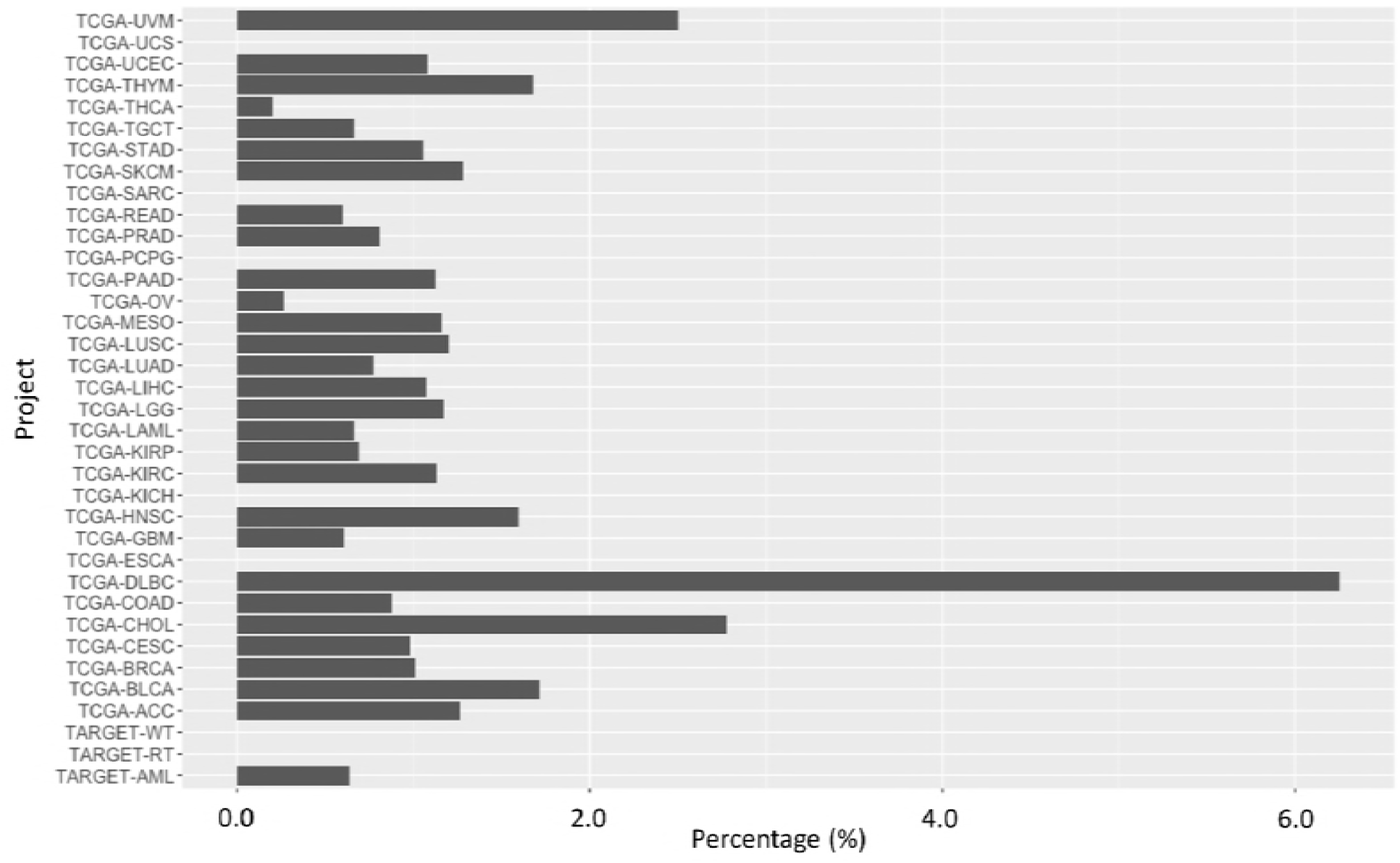
Percentage of cases that are ranked within top 100 in the predictive model over all cases in the TCGA project. The statistically significant overrepresented cancer type is B-cell lymphoma (TCGA-DLBC).

## Discussion

In this study, we computationally predicted and experimentally validated that levosimendan - a marked drug for heart failure – can inhibit the growth of multiple cancer cell lines, notably, lymphoma. The anti-cancer activity of levosimendan mainly origins from the modulation of RNA processing pathway by the inhibition of atypical kinase RIOK1. RIOK1 represents a new anticancer drug target[46, 47], and the chemical space of its inhibitors has just emerged. Tyrosine kinase inhibitors have been approved in the treatment of lymphoma[48, 49]. Our findings suggest that levosimendan could be used in the combination therapy or as a potential lead compound for new multi-targeted drugs for lymphoma. On one hand, levosimendan may be combined with other tyrosine kinase inhibitors that are associated with the risk of heart failure. Different from inhibitors generated from high-throughput screening or *de novo* design from a single target, it is known that levosimendan interacts with other proteins that are the drug targets for the heart failure than RIOK1. The combination of levosimendan and the tyrosine kinase inhibitor may not only reduce the cardiotoxicity of the tyrosine kinase inhibitor but also enhance the anti-cancer efficacy since they act on different cancer pathways. On the other hand, levosimendan interacts with multiple kinases that are associated with cancers as well as proteins that are responsible for heart failure. The binding promiscuity of levosimendan may allow us to use it as a lead compound to design a new type of dual action agent by modulating multiple targets that are involved in both side effects and disease mechanisms. In many cases, disease-causing genes have pleiotropic effects on biological system, thereby making on-target side effect(s) unavoidable. In contrast with the conventional drug discovery process that designs highly selective ligands, it is possible to mitigate the side effect by designing a drug to bind an off-target that is against the side effect[8].

To further advance the potential of levosimendan in cancer treatment, several questions remain to be answered. First, the anti-cancer potency of levosimendan can be further improved by designing personalized derivatives. The binding pose analysis may provide valuable clues to the drug design. Secondly, our machine learning model predicted that several other cell lines are sensitive to levosimendan. It will be interesting to verify these predictions. Finally, *in vivo* anticancer activity of levosimendan need to be verified.

The rational design of dual action multi-targeted drugs is an extremely challenging task. It requires modeling drug actions on a multi-scale, from genome-wide drug-target interactions to system level drug responses. This study showcases that 3D-REMAP is a potentially powerful tool towards designing polypharmacology and drug repurposing. 3D-REMAP provides a framework to integrate heterogeneous data from chemical genomics, structural genomics, and functional genomics, and synthesize diverse tools from bioinformatics, machine learning, biophysics, and systems biology for the multi-scale modeling of drug actions. Emerging paradigm of systems pharmacology enables the understanding of cellular mechanism of drug action at the organismal level, but lacks power to screen and design new chemical entities. Structure-based drug design has been successful in discovering novel drug molecules with fine-tuned binding properties to specific targets. However, the designed drug-target interaction may not transform into desired organismal level drug response. 3D-REMAP may bridge the structure-based drug design and systems pharmacology, thus it facilitates drug discovery for complex diseases. In spite of the success in this proof-of-concept study, many aspects of the 3D-REMAP platform could be improved. Firstly, the prediction accuracy of each individual algorithm such as protein-ligand docking and protein-chemical interaction prediction in the computational pipeline needs to be improved. Secondly, there is still a big gap between *in vitro* drug activity and clinical phenotype. More data types and modeling techniques, such as quantitative systems pharmacology and pharmacokinetics modeling as well as data mining of electronic health records, should be incorporated into the pipeline. With the knowledge of genome-wide target profiles and their disease associations, significant time and cost could be saved in the lead optimization, pharmacokinetics, pre-clinical, and other downstream studies using pro-target strategy for drug discovery.

## Material and Method

### 1. Materials

Solid levosimendan (Formula: CI_4_H_12_N_6_O, Molecular Weight: 280.28) was purchased from MedChemExpress (New Jersey, USA). The purity of the compound is larger than 98.0% determined by LCMS.

### 2. Kinase binding assay

To validate our computational predictions, we employed a competition binding assay to detect the binding of levosimendan to 425 human kinases. The proprietary KINOMEScan^™^ assay was performed by DiscoverX (Fremont, CA). The tests were performed at 10 μM and μM 100 concentrations of levosimendan, respectively. Assay results were reported as %Control, calculated as follows:

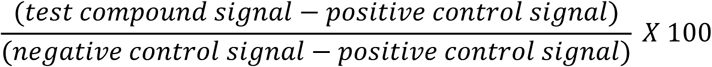

A lower %Control score indicates a stronger interaction. The KINOMEScan^™^ experiment and data analysis were performed by DiscoveRx (Fremont, CA).

### 3. Cancer cell line assay

OncoPanel^™^ cancer cell proliferation assay was performed by Eurofins Panlabs, Inc. (Missouri, USA). Cancer cells were grown in RPMI 1640, 10% FBS, 2 mM L-alanyl-L-glutamine, 1 mM Na pyruvate, or a special medium. Cells were seeded into 384-well plates and incubated in a humidified atmosphere of 5% CO2 at 37°C. Compounds were added the day following cell seeding. At the same time, a time zero untreated cell plate was generated. After a 3-day incubation period, cells were fixed and stained to allow fluorescence imaging of nuclei.

Levosimendan was serially diluted in half-log steps from 100 μM, and assayed over 10 concentrations with a maximum assay concentration of 0.1% DMSO. Automated fluorescence microscopy was carried out using a Molecular Devices ImageXpress Micro XL high-content imager, and images were collected with a 4X objective. 16-bit TIFF images were acquired and analyzed with MetaXpress 5.1.0.41 software.

Cellular response parameters were calculated using nonlinear regression to a sigmoidal single-site dose response model:

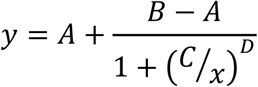

where *y* is the relative cell count measured following treatment with levosimendan at a concentration *x*. *A* and *B* are the lower and upper limits of the response, *C* is the concentration at the response midpoint (EC_50_), and *D* is the Hill Slope[50].

Cell count EC_50_ is the test concentration at the curve inflection point (parameter *C*), or half the effective response. IC_50_ is the test compound concentration at 50% of the maximum possible response. GI_50_ is the concentration needed to reduce the observed growth by half (midway between the relative cell count at the curve maximum and at the time of compound addition). Activity area is an estimate of the integrated area above the response curve[44]. Activity area values range from 0-10, where a value of zero indicates no inhibition of proliferation at all concentrations, and a value of 10 indicates complete inhibition of proliferation at all concentrations.

### 4. Computational methods

#### a. Ligand binding site similarity search across human structural proteome

The computational procedure has been reported previously[13-19]. Briefly, we used PDE3B (PDB ID 1SO2) that was the reported molecular target of levosimendan as the template for the binding site analysis. The SMAP software[30-32] was applied to characterize that protein’s ligand-binding potential from the geometric, physiochemical, and evolutionary characteristics of its binding pocket, and to predict the binding site similarity between the template and non-redundant human protein structures. The *p*-value of ligand binding site similarity was normalized by structural classes (all-alpha, all-beta, and mixed alpha-beta). The structures whose ligand binding sites were predicted to be similar to that of PDE3B with the *p*-value < 0.005 were selected as the initial candidate off-targets of levosimendan. Autodock Vina was used to predict the binding energy between selected off-targets and levosimendan, milrinone, anagrelide, amrinone, and enoximone. Drug-target interactions that had docking scores less than -7.5 were remained to be incorporated into the genome-wide chemical protein interactions network in the next step.

#### b. Genome-wide drug-target prediction

Genome-wide drug-target interactions were predicted using 3D-REMAP. The 3D-REMAP takes four networks as input: chemical-protein association, off-target, chemical-chemical similarity, protein-protein similarity networks, The chemical-protein associations were obtained by integrating three resources: 1) publicly available databases, ChEMBL[20] (v23.1) and DrugBank[21] (v5.5.10), 2) four data sets from recent publications about kinome assays[22-25], and 3) protein structure-based off-target prediction from previous step. From ChEMBL, inhibition assays having *IC*_50_ ≤ 10 *μM* was regarded as active associations. Those with suboptimal confidence scores (i.e. confidence < 9) were excluded. From DrugBank, drug-target, drug-enzyme, drug-carrier, and drug-transporter associations were collected. The data sets from kinome assays are available in different types of activity measurement. Christmann-Franck *et al*. collected chemical-kinase assays from multiple past publications and presented the activity standardization protocol, which assumed an activity with *K_i_* ≤*μM* is active[22]. If the original publication presented percent inhibition (or percent remaining activity) at a given compound concentration, *K_i_* was calculated as follows:

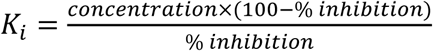

If the original publication presented *pK_i_* value, *K_i_* was obtained by *K_i_* = 10^−*pKi*^. For this study, we followed the above standardization protocol to integrate kinome assay data with the public databases. We considered chemical-kinase association active if *K_i_* ≤ 5 *μM* or *pK_i_* ≥ 5. To map chemicals from multiple sources, we used OpenBabel to convert all chemical molecules to InChIKey, a 27-character molecular representation developed to help searching chemical molecules. Protein targets were mapped by their UniProt accession. Low confidence targets from reference[24] were excluded. Off-target network was obtained using the procedure described in the previous section of Method. Chemical-chemical and protein-protein similarity scores were calculated similarly in the reference[26]. MadFast software developed by ChemAxon (https://chemaxon.com/) was used to calculate chemical-chemical similarity matrix, and BLAST was used to calculate protein-protein similarity matrix. The integrated chemical-protein association network contains 650,581 positively associated chemical-protein pairs for 1,656,274 unique chemicals and 9,685 unique target proteins. The chemical-chemical and protein-protein similarity matrices contain 122,421,717 and 31,266 nonzero similarity scores, respectively.

#### c. Gene expression and biological pathway analysis

Cell lines are classified as “sensitive” (activity area <= 1.95), “intermediate” (1.3 < activity area < 1.95), or “resistant” (activity area >= 1.3). Student’s t-test is used to compare log2-transformed mRNA probe levels between sensitive and resistant groups, and a p-value is calculated for each probe. The fold change in mRNA expression is calculated as:

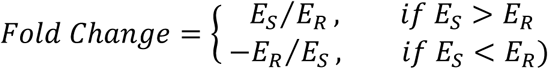

where *E_S_* and *E_R_* are the mean log2 mRNA probe levels for a given gene in cell lines found in the sensitive and resistant groups, respectively.

A false-discovery rate (FDR)-adjusted *p*-value (*q*-value) is computed with a null hypothesis of no difference between “sensitive” and “resistant” groups. The *q*-value is calculated according to the following formula:

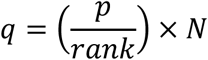

where *rank* is the rank of the p-value and *N* is the number of conducted tests.

Gene set over-representative analysis was carried out using DAVID[51].

#### d. Predictive modeling of levosimendan sensitivity of cancer cell lines and TCGA samples

The gene expression profiles of cell lines in CCLE[44] and GDSC[45] were used as features to build the pharmacogenomics model of levosimendan. Because of the inconsistency in the genomics data between these two data sets, two separate models were developed. 640 genes were identified to be responsible for the drug sensitivity (475 genes) and resistance (165 genes) based on the gene expression profile analysis as described in the previous section. They were used as initial gene set for the machine learning model. 97 and 85 genes were further selected via Kernel Conditional Covariance Mininization[43] for CCLE and GDSC data set, respectively. The gene expression profile of these genes were used to train and test the final models.

Using the gene expression profile of the selected genes as features and the activity area of cancer cell line sensitivity of levosimendan as the target variable, regression models were trained using ElasticNet, Random Forest, Support Vector Regression (SVR), and Gradient Boosting Regression as implemented in Scikit-learn. The features were standardized according to the machine learning algorithms applied. The optimal parameters were determined by grid search, and the performances were evaluated using nested leave-one-out cross-validation. ElasticNet and SVR were chosen as the best performed algorithms for CCLE and GDSC, respectively. After the models were trained, the response of remaining CCLE and GDSC cell lines that were not in the training data to levosimendan were predicted using corresponding models. The response of TCGA samples to levosimendan were predicted using CCLE trained model because the gene expression was measured by RNA-seq in both data sets.

#### e. Binding pose analysis

For experimentally validated kinase targets of levosimendan RIOK1, the binding pose of levosimendan was predicted using protein-ligand docking software AutodockFR[37] and visualized using DS Visualizer. Solved ADP-bound complex structure (PDB Id: 4OTP) was used for the docking experiment. Co-crystallized water molecules were remained in the docking.

## Acknowledgements

We greatly appreciate Drs. Alastair King and Charles Wageman (Eurofins Panlabs) for the valuable discussions.

## Author Contributions

H. L., Y. Q., D. H. performed experiments, analyzed the data, and wrote the manuscript; P. K. and X. S. performed experiments and analyzed the data; L.X. conceived the project, designed the experiments, analyzed the data, and wrote the manuscript.

